# Separating the genetic and environmental drivers of body temperature during the development of endothermy in an altricial bird

**DOI:** 10.1101/2024.06.21.600059

**Authors:** Lucy A. Winder, Jacob Hogger Gadsby, Eleanor Wellman, Joel L. Pick, Mirre J.P. Simons, Terry Burke

## Abstract

When altricial birds hatch, they are unable to regulate their own temperature, but by the time they fledge they are thermally independent. Early-life conditions have been shown to be an important factor contributing to an individuals’ performance in adult life. However, it is currently unknown to what extent body temperature during endothermy development is driven by genetic variation or by the early environment. We use thermal images of cross-fostered house sparrows (*Passer domesticus*) throughout the nestling period to separate genetic and environmental drivers of body temperature. Our results show small heritability of body temperature. We further found that there are effects from the natal environment which carry over into the late nestling stage. A correlation between early to the mid-nestling period was explained by the natal environment and during this period body temperature and growth follow independent developmental trajectories. Furthermore, higher body temperature is under viability selection independent of body mass. We, therefore, demonstrate the natal environment influences future offspring phenotype in a novel measure; body temperature. Our study provides the first study into the environmental and genetic drivers of body temperature variation in a wild bird, furthering our understanding of how an individuals’ traits evolve.

## Introduction

Passerine birds are heterothermic endotherms, whose body temperature is influenced by both intrinsic and extrinsic factors, such as body condition (Nord et al. 2013), time of day (Binkley et al. 1971; Barrett and Takahashi 1995) and exposure to stressors (Jerem et al. 2019). As adults, optimisation of body temperature has fundamental benefits for the individual, such as energy conservation (Tattersall et al. 2016) and improved ability to remain active (Torre-Bueno 1976). Altricial birds start to develop their own body temperature regulation as nestlings and complete this development before fledging. Nestlings are therefore sensitive to ambient temperature, which has been shown to impact their survival (Andreasson et al. 2018; Bourne et al. 2020; MarqueslJSantos and Dingemanse 2020). Previous studies on ambient temperature study its effects at the brood level, but we know little about how an individual’s body temperature is determined and this is yet to be considered in an evolutionary context.

Early life conditions affect the phenotype of developing individuals with effects persisting into adulthood (Metcalfe and Monaghan 2001; Saino et al. 2018; Spagopoulou et al. 2020). To date, the literature on body temperature in birds has largely focussed on the adaptive response to environmental conditions, mostly in experimental studies. Very few studies have been conducted in the wild and most of the literature on body temperature in birds comes from studies on poultry that have been selected for production. The rate of endothermy development in wild nestling blue tits (*Cyanistes caeruleus*) has been shown to vary between individuals and with brood size (Andreasson et al. 2016), and nestlings who have experimentally heated nests were better able to thermoregulate at later nestling ages (Page et al. 2022).

Early environments, such as the nest condition, may affect the thermal characteristics of juvenile birds, with future fitness implications. Indeed pre-hatching and post-hatching conditions have been shown to affect nestling phenotype, with growth a common trait measured, with maternal effects such as egg yolk composition and incubation behaviour also affecting nestling phenotype (Mousseau 1998; Saino et al. 2003; Nord and Nilsson 2011). However, we do not know what determines an individual’s development of body temperature regulation, and if the drivers of body temperature regulation are the same or separate from the drivers of growth.

Metabolic rate is a trait that is studied more commonly than body temperature but both probably share physiological mechanisms. Metabolic rate is a key trait for determining an individual’s energy expenditure (Biro and Stamps 2010). As body temperature is the result of metabolic heat production, this can be considered an indicator of an individual’s metabolic state. There is evidence that metabolic rate has a heritable component in captive zebra finches (Rønning et al. 2007) and wild blue tits, *Cyanistes caeruleus* (Nilsson et al. 2009). However, a recent study demonstrated that it is prenatal effects, rather than genetic effects, that drive metabolic rate variation in nestlings (McFarlane et al. 2021), which the previous studies were unable to disentangle. Metabolic rate has been shown to have an independent genetic basis from morphological and behavioural traits (Mathot et al. 2013) and also varies in how predictive it is of energy expenditure (e.g., Briga and Verhulst 2017). The study of body temperature can help us better understand how energy expenditure, growth and metabolic rate together determine physiological condition.

As a first step we aim to separate genetic and environmental effects on body temperature during development. There is some evidence in captive chickens that body surface temperature is heritable (Loyau et al. 2016). Altricial nestlings are easily cross-fostered between broods, and so provide the ideal opportunity to decompose variation in body temperatures into genetic, pre- and post-natal parental (or nest) effects (Winney et al. 2015). Here, we first decompose the variation in body temperature through ontogeny into genetic, natal environment and rearing environment effects, covering a period of pre- and post-endothermy development. We also determine, at a range of nestling ages, whether body temperature is under selection independently of growth. Our study shows that early-life conditions influence an individual’s phenotypes using a non-invasive measure of individual quality, body surface temperature.

## Materials and methods

### Study population

Data for this study comes from a long-term study of a closed population of house sparrows (*Passer domesticus)* on Lundy Island in the Bristol Channel, (51°100 N, 4° 400 W), monitored since 1996. LAW collected the thermal data for this study during the breeding season in 2018 and 2019. LAW cross-fostered nestlings in 2018 and 2019 at two days of age post-hatching (day first chick observed in nest = day 1). Where possible, we used a triad approach, where up to three broods matched by their hatching date had nestlings swapped between them. As many nestlings were cross-fostered as possible – broods of equal size had all nestlings cross-fostered and brood of differing in size had the number of chicks in the smallest clutch cross-fostered. If only one brood was at age two on a given day, then no cross-fostering occurred. 16 nestlings were removed from the dataset as they hatched significantly later than their siblings of the same brood (nestlings in a given brood generally hatch within 24 hours of each other). Parents of the focal birds were identified from unique colour ring combinations and a British Trust for Ornithology metal ring. A genetic pedigree was assembled using 13 microsatellite loci from blood samples (Dawson et al. 2012 as described in Schroeder et al. 2015). All blood sampling procedures were performed under a UK Home Office licence.

### Thermal image collection

Surface temperature has previously been shown to correlate linearly with core temperature in birds (Hill et al. 1980; Giloh et al. 2012). The use of non-invasive temperature measurements via thermal imaging of surface temperature can therefore be used to determine an individual’s thermal state (McCafferty 2013; Nord et al. 2016). Eye-region temperature is the hottest surface region of passerine birds (Jerem et al. 2015). There is also evidence that eye-region temperature remains relatively stable when birds are faced with a perceived risk of an energetic shortfall, suggesting the eye region offers a reliable surface correlate of core temperature (Winder et al. 2020). Here, we use eye-region temperature (or maximum head temperature in very young birds, see below) measurements throughout the nestling period to determine to what extent body temperature is driven by natal and rearing conditions.

LAW visited each hatched brood on four occasions, when the nestlings were 2 (n= 696 individuals, 231 broods), 5 (n= 582 individuals, 216 broods), 10 (n= 519 individuals, 201 broods) and 12 (n= 490 individuals, 191 broods) days old. House sparrow nestlings have been shown to develop endothermy by 9.5 days post-hatching (Dunn 1975). Therefore, this age range captures a near-poikilothermic (i.e., body temperature which varies with the environment) stage through to that of thermal independence. We limited our visits to the morning to minimise the effect of time of day. On each visit, LAW obtained a thermal image of each nestling, as follows. We temporarily removed all nestlings from the nest to allow measurement. LAW then took a thermal image of the right side of each bird’s head using a C3 FLIR camera (FLIR® Systems, Inc.). Birds within a nest were thermally photographed in a randomised order, though order did not affect body temperature readings (data not shown) and so was not included in the final statistical models described below. The air temperature and relative humidity were also recorded at the time of image capture. Individual birds were identified from a toenail clipped on day 2.

### Temperature extraction

JHG and EW extracted the nestling maximum head temperature (hereafter referred to as body temperature, for simplicity) from each image using the Thermimage package (Tattersall 2017) in R version 3.6.2 (R Development Core Team 2009) and by selecting the hottest pixel in the image of the bird’s head. The hottest pixel was largely on the eye region of the bird, however, when birds are very young, tissue and organs are not differentiated in our thermal images and so the hottest pixel on the bird’s head was used. In some cases, the hottest pixel was not on the bird but instead in the background of the image (e.g., when birds were in poor health and colder than the environment) and, therefore, JHG and EW extracted the maximum head temperature identified in FLIR Tools by drawing a temperature selection box exclusively around the bird’s head. The atmospheric temperature and relative humidity (from recordings taken at the time of measurement) were corrected for during image extraction in either Thermimage or FLIR Tools.

### Data analysis

LAW performed all statistical analyses for this study with guidance from JLP, who also reviewed the analysis code. LAW determined the heritability of body temperature for each age (2, 5, 10 and 12 days) in separate “animal models”, which use the relatedness structure of a pedigree to estimate additive genetic effects (Henderson 1988; Kruuk 2004). We included fixed effects of air temperature (continuous), year (2 level factor) and brood size (at given age; continuous) in the models. In addition to additive genetic effects, the natal brood (brood in which the nestling was born into) and the rearing brood (brood in which the nestling was raised) were modelled as random effects. Random effects of rearing brood were not included in the model for day 2 as temperature measures were taken at the point of cross-fostering. All covariates were mean-centred to aid interpretation.

To account for selective disappearance when estimating the variance components of body temperature (i.e., there is a potential to underestimate the influence a trait has on viability when an individual with a given phenotype dies before it is measured at a later age), LAW then modelled body temperature (for all ages combined) using a multivariate linear mixed effects model (Hadfield 2008; Hadfield et al. 2013). Fixed effects of age, brood size (at given age), air temperature and year were modelled separately for each age. We modelled natal brood effects with a 4*4 unstructured covariance matrix, and the same covariance structure for the residual effects. We modelled rearing brood effects for ages 5, 10 and 12 with a 3*3 unstructured covariance matrix. We did not have sufficient power to model genetic effects in this analysis, most likely due to the very low genetic variance across ages. When natal brood is separated from the rearing brood in our models, the genetic variance will be confounded with the natal brood effects; however, given the small additive genetic effects in the models of each age separately this will be negligible (at least at older ages). We repeated this analysis but included a fixed effect of body mass (again separately for each age) to determine to what extent body temperature is a separate trait from body mass.

The analyses described above were run in MCMCglmm version 2.36 (Hadfield 2010) using R version 4.4.0 (R Development Core Team 2009). The burn-in period was 45,000 for all analyses and a chain length of 195,000 iterations and a thinning interval of 150. Parameter expanded priors were used for the random effect variances (V=1, nu=1, alpha.mu=0 and alpha.V=1000) and inverse wishart priors were used for the residual variances (V = 1 and nu = 0.002 for the univariate models, and V= 1e-6 and nu=5 for the multivariate models). A p-value for the fixed effects and covariances in these models was approximated (pMCMC) as two times the smaller number of iterations where the parameter value is either less than zero or greater than zero (Baayen et al. 2008), equivalent to a two-sided test. When model estimates are presented, we use the posterior median (unless otherwise stated).

LAW analysed whether body temperature predicts survival to the next age using generalised linear mixed effect models with a binomial error distribution in lme4 (Bates et al. 2015), using the bobyqa optimiser. For age 12, survival was determined by whether an individual was sighted as an adult (i.e., over winter or during the following breeding season). All ages were analysed in separate models, and included fixed effects of body temperature, body mass and year, and random effects of natal brood (all ages) and rearing brood (Age 5, 10 and 12 only).

## Results

### Variance explaining body temperature at each age

To reliably estimate the environmental and genetic variance components explaining body temperature at each age, we accounted for air temperature, which as expected, was significantly positively correlated with body temperature at all ages (Table 1). As our study took place over two consecutive years, we also accounted for differences in the environmental conditions between years, by incuding it as a two-level factor in the model. In 2019, birds were warmer at ages 2 and 5 compared to those recorded in 2018. However, birds were cooler at ages 10 (not significant) and 12 compared to 2018 (Table 1).

**Table 1:**
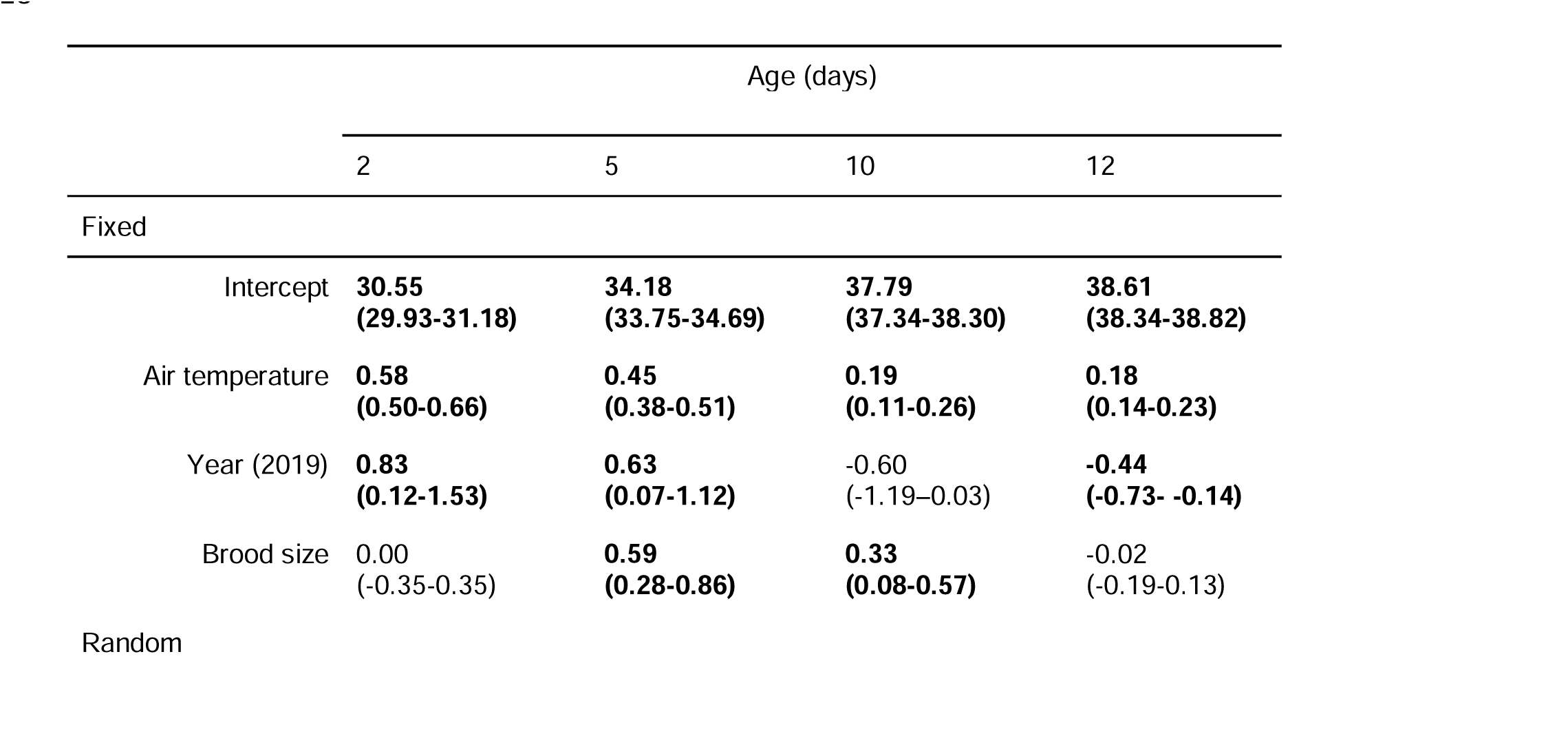
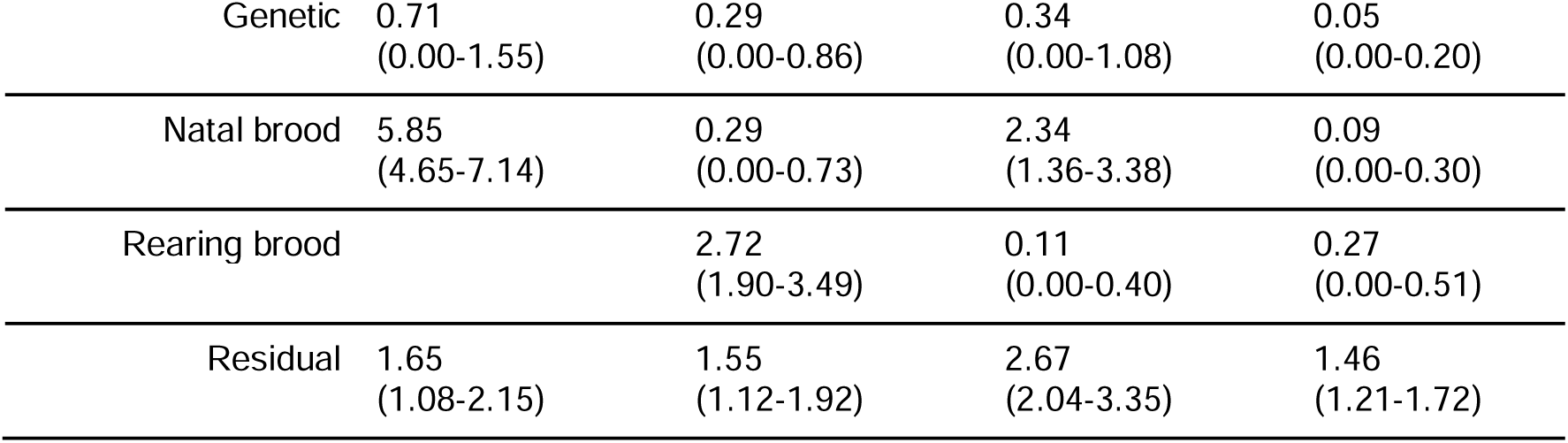
Variance components of body temperature by age. Values for fixed and random effects are the posterior mean (95% credible intervals). Statistically significant values (pMCMC <0.05) are in bold.

|  |  | Age (days) |  |  |  |
| --- | --- | --- | --- | --- | --- |
|  |  | 2 | 5 | 10 | 12 |
| Fixed |  |  |  |  |  |
| Intercept |  | **30.55**<br>(29.93-31.18) | **34.18**<br>(33.75-34.69) | **37.79**<br>(37.34-38.30) | **38.61**<br>(38.34-38.82) |
| Air temperature |  | **0.58**<br>(0.50-0.66) | **0.45**<br>(0.38-0.51) | **0.19**<br>(0.11-0.26) | **0.18**<br>(0.14-0.23) |
| Year (2019) |  | **0.83**<br>(0.12-1.53) | **0.63**<br>(0.07-1.12) | -0.60<br>(-1.19-0.03) | **-0.44**<br>(-0.73- -0.14) |
| Brood size |  | 0.00<br>(-0.35-0.35) | **0.59**<br>(0.28-0.86) | **0.33**<br>(0.08-0.57) | -0.02<br>(-0.19-0.13) |
| Random |  |  |  |  |  |

|  |  |  |  |  |
| --- | --- | --- | --- | --- |
| Genetic | 0.71<br>(0.00-1.55) | 0.29<br>(0.00-0.86) | 0.34<br>(0.00-1.08) | 0.05<br>(0.00-0.20) |
| Natal brood | 5.85<br>(4.65-7.14) | 0.29<br>(0.00-0.73) | 2.34<br>(1.36-3.38) | 0.09<br>(0.00-0.30) |
| Rearing brood |  | 2.72<br>(1.90-3.49) | 0.11<br>(0.00-0.40) | 0.27<br>(0.00-0.51) |
| Residual | 1.65<br>(1.08-2.15) | 1.55<br>(1.12-1.92) | 2.67<br>(2.04-3.35) | 1.46<br>(1.21-1.72) |

We found that body temperature had a small heritability at age 2 and reduced as the nestlings aged (Age 2, h^2^ = 0.08 (95% CI: 0.00-0.21); Age 5, h^2^ = 0.04 (95% CI: 0.00-0.23); Age 10, h^2^ = 0.03 (95% CI: 0.00-0.23); Age12, h^2^ = 0.01 (95% CI: 0.00-0.13); Figure 1, Table 1). At early ages (age 2 and 5), body temperature was driven by the current conditions the individual experiences in the nest (i.e., the combined natal and rearing brood effect for age 2 and rearing brood for age 5). At age 10, when the nestlings become thermally independent, temperature was not driven by their current environment (rearing) but by their prior natal environment. The amount of residual variation in body temperature increased as the nestlings aged, suggesting that environmental variables not captured in our models become increasingly determinative of body temperature. By age 12, only a small proportion of variation was driven by the rearing brood (and negligible variation was driven by genetic or the natal environment) and most variation was explained by residual variation (Table 1, Figure 1).

**Figure 1:**
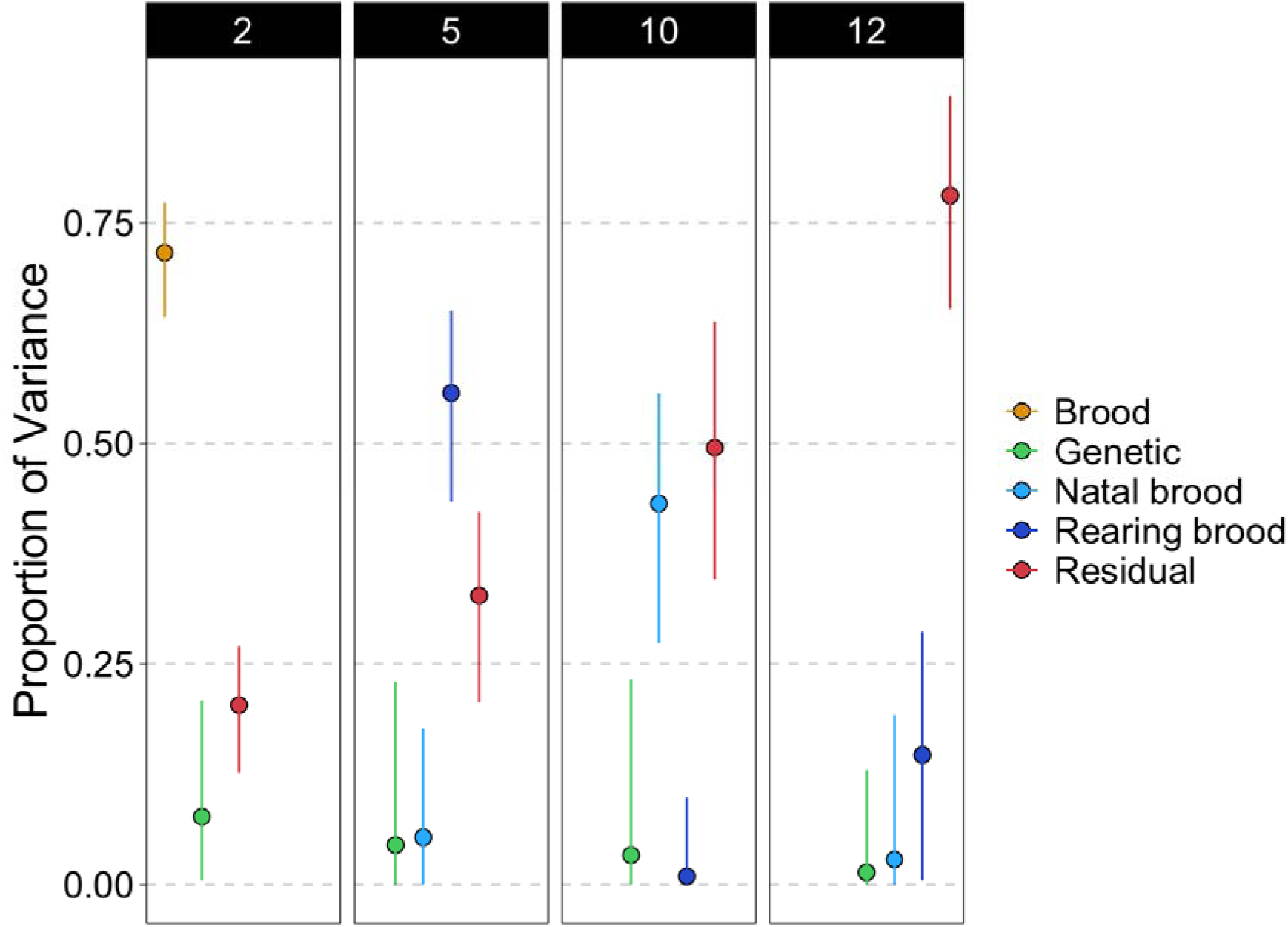
The proportion of variance in body temperature at each age (2, 5, 10 and 12 days post-hatching) over the nestling period for the random effects in the model. Note birds were cross-fostered at age 2 and brood is the combination of rearing and natal brood (which are both the same at this age).

### Covariance in body temperature across ages

We found that chicks that hatched in the same nest who have higher temperatures just after hatching (2 days) also have higher temperatures at later ages (5 and 10 days) (Table 2).

**Table 2:** Posterior medians (95% credible intervals) for the random effects of the model of body temperature by age. Variances (95% credible intervals) are on the diagonal. Covariances are below the diagonal and correlations (95% credible intervals) are above the diagonal. Significant medians and correlations are in bold.

|  | 2 | 5 | 10 | 12 |
| --- | --- | --- | --- | --- |
| Natal Brood: |  |  |  |  |
| 2 | 6.05<br>(4.94, 7.51) | **0.56**<br>**(0.22, 0.97)** | **0.31**<br>**(0.12, 0.48)** | -0.03<br>(-0.88, 0.80) |
| 5 | **0.85**<br>**(0.25, 1.48)** | 0.38<br>(0.06, 0.96) | 0.50<br>(-0.06, 0.93) | -0.29<br>(-0.99, 0.99) |
| 10 | **1.15**<br>**(0.41, 1.98)** | 0.45<br>(-0.05, 1.13) | 2.42<br>(1.52, 3.50) | 0.11<br>(-0.82, 0.82) |
| 12 | -0.03<br>(-0.45, 0.37) | -0.05<br>(-0.35, 0.16) | 0.05<br>(-0.32, 0.59) | 0.25<br>(0.00, 0.70) |
| Rearing Brood: |  |  |  |  |
| 2 | / | / | / | / |
| 5 | / | 2.63<br>(1.94, 3.52) | 0.68<br>(-0.47, 0.99) | 0.30<br>(-0.70, 0.97) |
| 10 | / | 0.39<br>(-0.05, 1.03) | 0.14<br>(0.00, 0.64) | 0.42<br>(-0.84, 0.98) |
| 12 | / | 0.17<br>(-0.19, 0.55) | 0.03<br>(-0.08, 0.26) | 0.15<br>(0.00, 0.53) |
| Residual: |  |  |  |  |
| 2 | 2.06<br>(1.82, 2.325) | **0.15**<br>**(0.05, 0.25)** | 0.07<br>(-0.04, 0.18) | -0.05<br>(-0.16, 0.06) |
| 5 | **0.28**<br>**(0.09, 0.47)** | 1.72<br>(1.49, 2.00) | **0.14**<br>**(0.02, 0.25)** | 0.09<br>(-0.05, 0.23) |
| 10 | 0.17<br>(-0.09, 0.43) | **0.31**<br>**(0.05, 0.57)** | 2.83<br>(2.38, 3.44) | **0.15**<br>**(0.01, 0.28)** |
| 12 | -0.10<br>(-0.32, 0.11) | 0.16<br>(-0.08, 0.42) | **0.33**<br>**(0.03, 0.68)** | 1.87<br>(1.55, 2.28) |

This relationship weakened slightly when body mass was included in the model, but remained significant between ages 2 to 10 (Table 3). However, by age 12 the correlation with early ages disappears (Table 2 & 3; although note the large confidence intervals for these estimates, likely driven by the low variances in this component at this age). We also found that nestlings raised together were no more likely to have similar temperatures at later ages (Table 2 & 3), suggesting that body temperature is an individual-based trait.

**Table 3:**
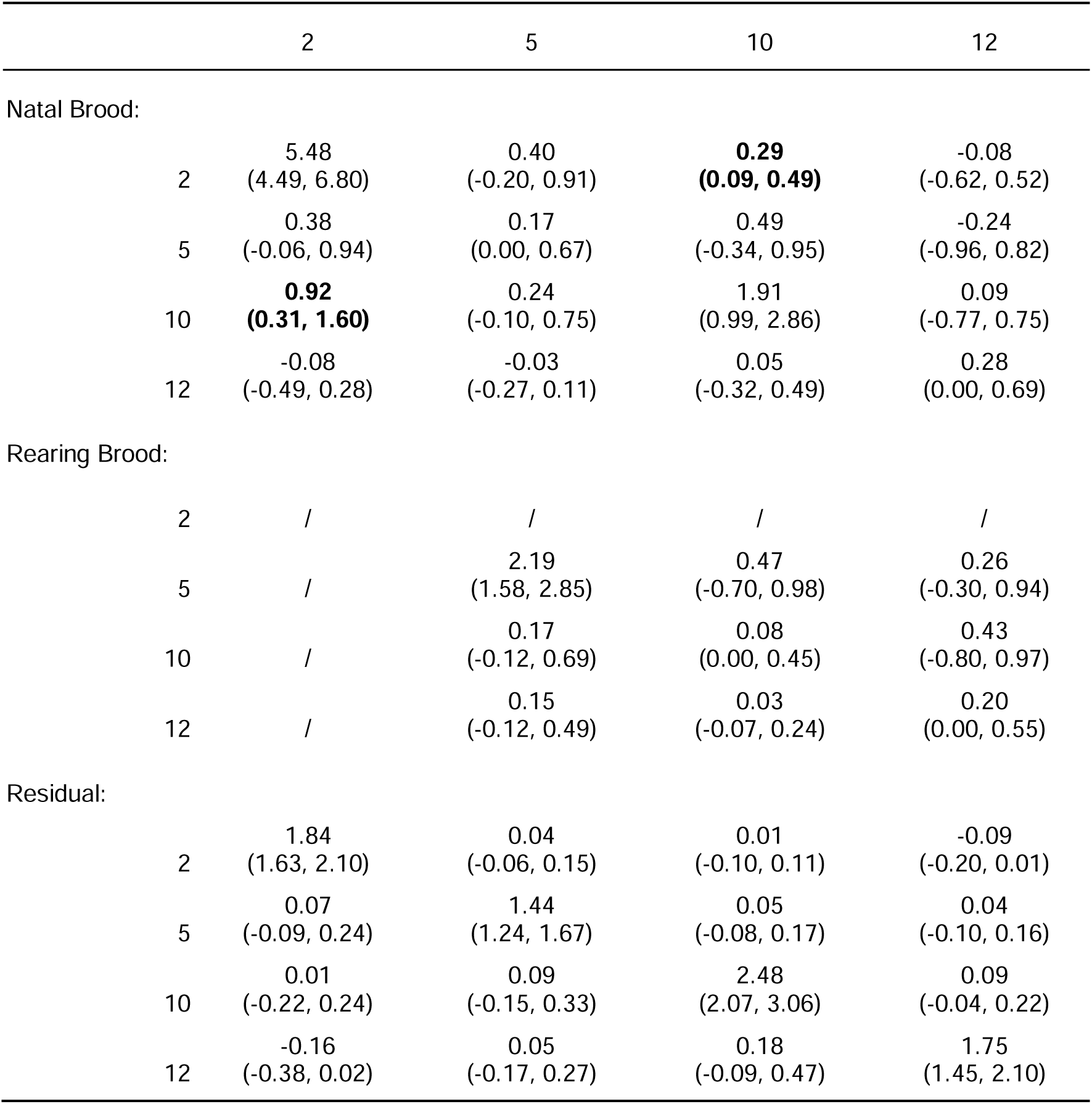
Posterior medians (95% credible intervals) for the random effects of the model of body temperature by age. The model included an interaction term between age and body mass. Variances (95% credible intervals) are on the diagonal. Covariances are below the diagonal and correlations (95% credible intervals) are above the diagonal. Significant medians and correlations are in bold.

### Selection on body temperature

Nestling body temperature had a positive effect on their survival to the next age, independent of body mass (Table 4; although this was only statistically significant at ages 2 and 10). However, body temperature at age 12 did not predict survival to adulthood, though only 12 (out of 490) individuals survived to adulthood (estimate = 0.00 ± 0.04, Table 4). Body mass also predicted survival but only at ages 2 and 5 (Table 4).

**Table 4:** Model outputs for survival to the next age class predicted by body temperature at each age. Survival for age 12 was determined by being resighted as an adult.

|  |  | Estimate | SE | z | p |
| --- | --- | --- | --- | --- | --- |
| Age 2 |  |  |  |  |  |
| *Fixed* |  |  |  |  |  |
|  | (Intercept) | 0.99 | 0.35 | 2.86 | **0.004** |
|  | Body temperature | 0.31 | 0.07 | 4.25 | **<0.001** |
|  | Body mass | 0.91 | 0.19 | 4.83 | **<0.001** |
|  | Year (2019) | 3.49 | 0.66 | 5.26 | **<0.001** |
| *Random* |  |  |  |  |  |
|  |  | Variance | SD |  |  |
|  | Natal brood | 8.88 | 2.98 |  |  |
| Age 5 |  |  |  |  |  |
| *Fixed* |  |  |  |  |  |
|  | (Intercept) | 1.38 | 0.40 | 3.47 | **<0.001** |
|  | Body temperature | 0.18 | 0.10 | 1.88 | 0.060 |
|  | Body mass | 0.49 | 0.10 | 4.77 | **<0.001** |
|  | Year (2019) | 3.65 | 0.70 | 5.23 | **<0.001** |
| *Random* |  |  |  |  |  |
|  |  | Variance | SD |  |  |
|  | Natal brood | 0.36 | 0.60 |  |  |
|  | Rearing brood | 7.22 | 2.69 |  |  |
| Age 10 |  |  |  |  |  |
| *Fixed* |  |  |  |  |  |
|  | (Intercept) | 6.58 | 1.19 | 5.51 | **<0.001** |
|  | Body temperature | 0.72 | 0.22 | 3.32 | **<0.001** |
|  | Body mass | 0.11 | 0.09 | 1.25 | 0.21 |
|  | Year (2019) | 5.17 | 1.90 | 2.72 | **0.007** |
| *Random* |  |  |  |  |  |
|  |  | Variance | SD |  |  |
|  | Natal brood | 9.40 | 3.07 |  |  |
|  | Rearing brood | 121.34 | 11.02 |  |  |
| Age 12 |  |  |  |  |  |
| *Fixed* |  |  |  |  |  |
|  | (Intercept) | -7.74 | 1.29 | -6.00 | **<0.001** |
|  | Body temperature | -0.29 | 0.38 | -0.76 | 0.45 |
|  | Body mass | 0.10 | 0.14 | 0.72 | 0.47 |
|  | Year (2019) | -14.51 | 418.05 | -0.04 | 0.97 |
| *Random* |  |  |  |  |  |
|  |  | Variance | SD |  |  |
|  | Natal brood | 0.00 | 0.00 |  |  |
|  | Rearing brood | 37.43 | 6.12 |  |  |

## Discussion

Our study provides a crucial step in the understanding of the drivers of a key physiological trait in birds, body temperature. We found evidence of heritability of body temperature, though this reduced as the nestlings aged. Our results also show that the natal brood explains a significant amount of variance in body temperature after being cross-fostered and after endothermy developed. A notable finding of this work was that variation in body temperature and its effect on survival were, to a degree, separate from growth. Our results provide a crucial step in understanding the drivers of a key physiological trait in birds, body temperature.

A key finding of this work is that early life effects predict body temperature in later nestling ages. The combination of small or negligible heritability along with variation explained by the natal brood provides evidence that the pre-hatching effects have a profound impact on the physiology of nestlings during their development. Possible mechanisms for this could be either parental incubation effects or investment in egg quality (for example, through investment of proteins or hormones). Environmental conditions during development have been shown to affect nestling physiology, morphology and survival (Nord and Nilsson 2011; Ospina et al. 2018; Mueller et al. 2019). However, nestlings can compensate for negative early life conditions (Metcalfe and Monaghan 2001; Bize et al. 2006). This could, in part, explain why, by age 12, variance explained by the natal and rearing environments was low (though see below where we discuss selective disappearance). At age 10, after endothermy has developed, the natal brood explained more variation in body temperature than the current environment, the rearing brood. This was the case when body temperature was modelled for each age individually and additionally when selective disappearance was accounted for, where body temperature was largely explained by covariances associated with natal brood effects between ages 2 to 5 and 2 to 10.

However, when body mass, as well as selective disappearance, was accounted for, early-stage body temperature did correlate with mid-stage nestling temperature (i.e., 2 to 10 only) but the effects of natal and rearing brood could not be separated for these ages. This does still importantly show that early life effects have significant carry-over effects to the later stages of the nestling period and that this occurs separately from the effects of growth, though further work is needed to determine if it is the natal parents or social parents that drive this variation. Intuitively, Thomson et al. (2017) found a small effect of the nest of origin and heritability of body mass and found as blue tit (*Cyanistes caeruleus*) nestlings age, body mass becomes more driven by the social parents. However, our results do not provide evidence for the same trend as rearing brood explains little variation in any of our models, though this variation is confounded with that of the natal brood for the age 2 effects.

Our findings also importantly show that the drivers of body temperature are fundamentally different to those of body mass, despite being correlated (Supplementary information S1). Previous studies have shown that nest temperature affects survival (Berntsen and Bech 2016; Andreasson et al. 2018). Our study, importantly, goes beyond the collective nest effect and explores body temperature at the individual level. Determining the survival effects of body temperature at the individual level also allowed us to separate this effect from other physiological effects that may impact survival. We found that selection is acting on body temperature; body temperature predicts survival to the next age, with warmer birds being more likely to survive. The selection on body temperature was independent of body mass, showing the selection effect of body temperature is not simply a result of larger nestlings being selected for. Interestingly, at age 10, body mass did not significantly predict survival, but body temperature did. This demonstrates the potential for body temperature to be a key indicator of individual quality during the nestling period. By age 12, this trend disappeared, possibly because by this age, the cooler, poor-quality nestlings had died (i.e., selective disappearance). It is also possible that, after fledging, many other extrinsic factors lead to the death of an individual and therefore, the body temperature is negligible in explaining mortality. Indeed, very few individuals survived to adulthood and so the results of survival beyond fledging should be treated with caution. Body temperature therefore indicates the quality of a nestling that is independent of growth, although this may not translate into adulthood. It would also be interesting, though challenging in terms of the sample size required, to follow these effects throughout adulthood to determine if there are subsequent fitness consequences of nestling body temperature.

## Conclusion

Our results from this study provide insights into what drives variation in body temperature throughout the nestling period and that body temperature is somewhat independent of the effects of growth. This study also demonstrates the importance of determining what drives variation in phenotypic traits in early life, as without cross-fostering, it would be difficult to separate rearing effects from pre-hatching incubation effects. Importantly, our study shows the rearing environment explains very little variation in body temperature after endothermy has developed. We have also shown that body temperature is selected for independently of body mass, and therefore, body temperature could provide a useful tool for determining differences in individual quality.

## Acknowledgements

Author initials are provided as per the MeRIT system (Nakagawa et al. 2023). MJPS is funded by a Sir Henry Dale Fellowship, Wellcome and Royal Society (216405/Z/19/Z).

