## Supplementary Information for "Separating the genetic and environmental drivers of body temperature during the development of endothermy in an altricial bird"

S1: The correlation between body mass and body temperature at each age. Regression lines are from locally estimated scatterplot smoothing.

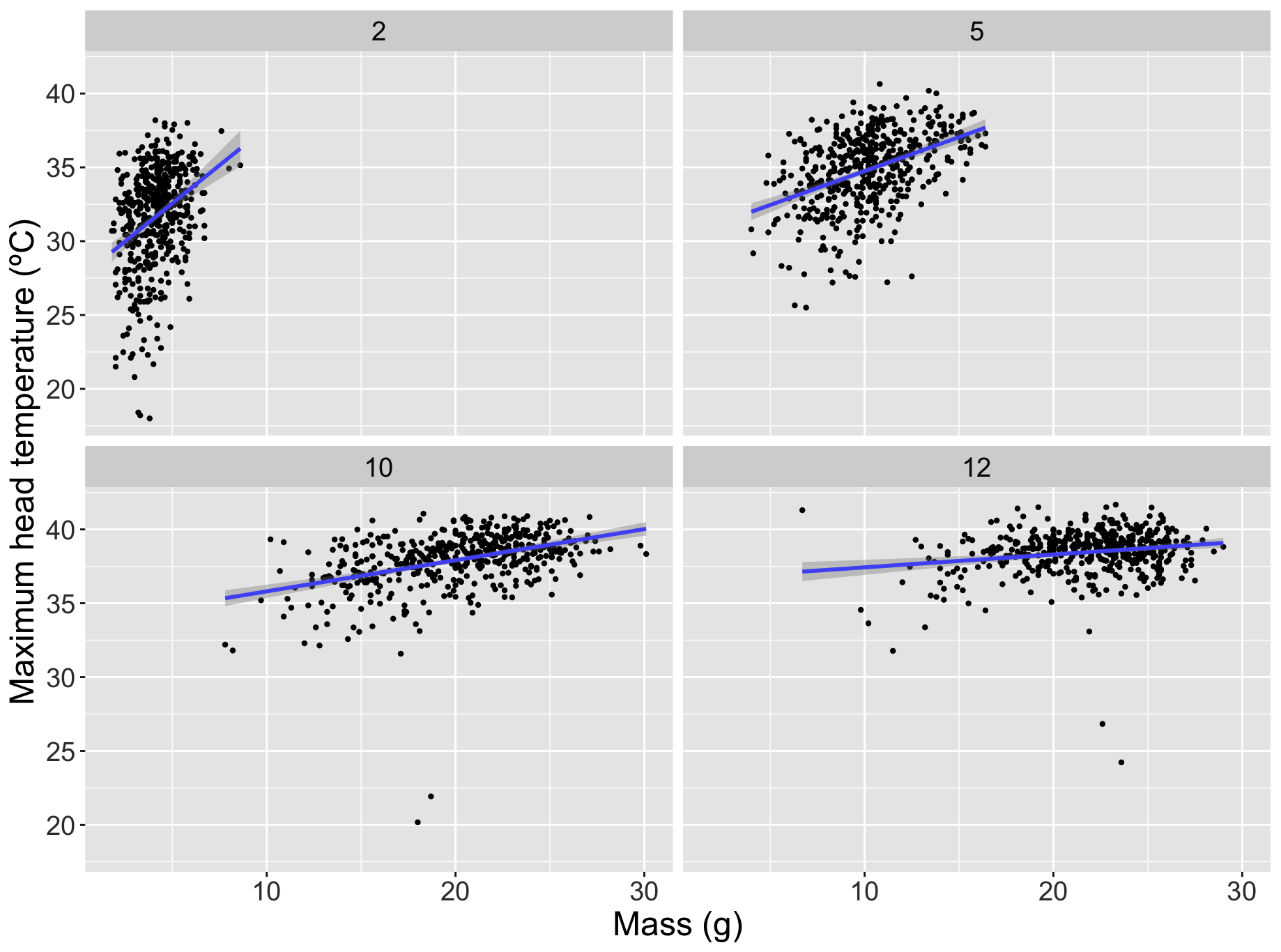
